# OPETH: Open Source Solution for Real-time Peri-event Time Histogram Based on Open Ephys

**DOI:** 10.1101/783688

**Authors:** András Széll, Sergio Martínez-Bellver, Panna Hegedüs, Balázs Hangya

## Abstract

Single cell electrophysiology remains one of the most widely used approaches of systems neuroscience. Real-time feedback during electrophysiology experiments is important to guide experimental decisions that eventually determine the quality of recording, duration of the project and value of the collected data. We present an open-source tool that enables flexible online visualization of action potential alignment to external events, called the peri-event or peristimulus time histogram (OPETH). Based on the Open Ephys open source data acquisition system, we developed a Python interface for real-time plotting of neuronal spike times and evoked waveforms with respect to external events represented by digital logic signals. These digital inputs may signal photostimulation time stamps for *in vivo* optogenetic identification of cell types or the times of behaviorally relevant events during *in vivo* behavioral neurophysiology experiments. Therefore, OPETH allows real-time identification of genetically defined neuron types or behaviorally responsive populations. By allowing ‘hunting’ for neurons of interest, OPETH may significantly increase the efficiency of experiments that combine *in vivo* electrophysiology with behavior or optogenetic tagging of neurons.

## Introduction

Neurons are diverse. Often they are categorized based on their neurotransmitters, neuropeptides, calcium-binding proteins, ion channels and other markers (Ascoli et al., 2008; Klausberger and Somogyi, 2008). These usually entail the specific expression of proteins, which provides a genetic handle on these cell types (Harris et al., 2018). This allowed the recent introduction of optogenetic cell type identification or optogenetic tagging (Cohen et al., 2012; Kvitsiani et al., 2013; Lima et al., 2009). In brief, the expression of a restriction endonuclease, most often Cre, is controlled by a genomic promoter, and a light sensitive ion channel or pump is introduced in a Cre-dependent manner (Boyden et al., 2005). Thus, the cell type defined by the promoter becomes photosensitive, allowing their identification on extracellular recording: the cells that respond to light with short latency, precisely timed action potentials belong to the given class (Hangya et al., 2014; Kvitsiani et al., 2013; Lima et al., 2009). Previously, *in vivo* cell type identification was only possible with juxtacellular recording and labeling or *in vivo* whole cell patch clamp, which were mostly restricted to anesthetized recordings (Pinault, 1996). While still often relatively low yield, optogenetic tagging opened the gate for larger scale recording of identified neurons in awake behaving animals. However, neurons are often identified during offline analysis, which limits the flexibility and planning of the experiments, resulting in lower number of tagged cells and longer projects.

A caveat of optogenetic tagging studies is that light may induced different signals besides action potentials, including photoelectric (Becquerel) and photovoltaic effects (Kozai and Vazquez, 2015), or exciting too many neuronal elements summing up to population spikes that prevent proper spike sorting. The uneven dispersion of light in brain tissue may lead to artifacts that are hard to remove by offline referencing techniques, as pointed out in previous studies (Cardin et al., 2010; Mikulovic et al., 2016; Park et al., 2014). Most of these potential confounds can be efficiently eliminated by proper control of light intensities delivered into the brain, for which precise on-line feedback is immensely useful.

Additionally to genetically defined types, neurons are often characterized by the relation of their firing pattern to external events *in vivo*. For instance, neurons of sensory cortices are categorized by their response to sensory stimuli (Gentet et al., 2012; Hires et al., 2015); in reverse, the features of sensory events that activate a given neuron gave rise to the concept of the receptive field (Hubel and Wiesel, 1959; Kilgard and Merzenich, 1998; Ko et al., 2011). Neurons thought to participate in cognitive processing are analyzed with respect to the salience and motivational value of external stimuli (Hangya et al., 2015; Lin and Nicolelis, 2008; Schultz et al., 1997), while neurons on the effector side are correlated with muscle activity and movements (DeLong, 1971). To visualize and quantify the correlation between these external events and neural activity, a linear correlation technique called the peri-event or peri-stimulus time histogram is usually applied (Endres et al., 2008; Solari et al., 2018). The PETH is a histogram of relative action potential times with respect to the event of interest; thus, it is mathematically equivalent to the cross-correlation (convolution) of spike and event times. When aiming to study a specific group of neurons, e.g. classically tuned neurons of the primary auditory cortex (Hromádka et al., 2008; Pi et al., 2013), or reward activated neurons of the VTA (Cohen et al., 2012; Schultz et al., 1997), it is particularly helpful to have a real-time PETH readout during positioning the recording electrodes.

Therefore we developed a real-time ‘online’ PETH or OPETH based on the Open Ephys open source data acquisition system (Siegle et al., 2017). This constitutes of a modified ZeroMQ plugin for distributed messaging and a Python interface that receives the data and visualizes peri-event time histograms and evoked waveforms quasi real time. These tools are useful for tracking the neuronal responses to light stimulation for optogenetic tagging or to behaviorally relevant events during animal training. We find that OPETH helps guide experimental decisions and greatly speeds up optogenetic experiments by allowing ‘hunting’ for tagged neurons.

## Methods

In this section we provide a system overview, a description of the Open Ephys ZMQ plugin and setting up the signal chain and the detailed presentation of the Python GUI interface for OPETH. Source code is available at https://github.com/hangyabalazs/ZMQInterface.git and https://github.com/hangyabalazs/opeth.git.

### System overview

Animals were implanted with custom-built implants that include Omnetics connectors that can interface with the Intan RHD2000 chip series, compatible with the Open Ephys system (Siegle et al., 2017; Solari et al., 2018). Data were amplified, digitized and digitally multiplexed by one or two 32-channel Intan Headstages RHD2132, providing 32- or 64-channel digital recordings. Data were transferred to the Open Ephys acquisition board by Intan Serial Peripheral Interface (SPI) cables.

Neural data were acquired at 30 kHz sampling rate by the open source, plug-in based Open Ephys Data Acqusition System. We used a modified ZMQ plug-in (https://neuroinformatics.nl/drupal/?q=node/181) to stream data to external programs and accessed the ZMQ data stream from Python programming language. The OPETH GUI, implemented in Python, visualizes online PETH and evoked waveform plots, providing access to spike discrimination thresholds and other parameters.

We used the BPod Behavior Control System (Sanworks Inc.) for real-time behavioral control during animal training. BPod is an open source, microcontroller-based system implementing a finite state machine optimized for low latencies that allow the combination of electrophysiology, optogenetics and animal behavior (https://sanworks.io/shop/products.php?productFamily=bpod). BPod sent TTL pulses at each stimulus onset and reward (water) or punishment (air puff) delivery to synchronize behavioral events with neural recordings.

We used the PulsePal stimulator (Sanworks Inc.) to trigger 1 ms square pulses of a blue laser (Sanctity Laser, SSL-473-0100-10TM-D-LED) at 20 Hz with 2 s ON - 3 s OFF duty cycle. The laser light was delivered to the target area by a patch cable (Thorlabs), LC-LC type optical connectors (Thorlabs) and a 50µm core optical fiber (Laser Components) for optogenetic tagging. TTL pulses were sent both to the blue laser and to Open Ephys to synchronize photostimulation and recording.

System components are summarized in Table 1.

**Table 1:** List of main components used during the experiments

| Device | Company |
| --- | --- |
| Intan Headstage RHD2132 | Intan Technologies |
| Serial Peripheral Interface cables | Intan Technologies |
| Open Ephys acquisition board | Open Ephys |
| Open Ephys Input Output board | Open Ephys |
| BPod Behavior Control System | Sanwork Inc. |
| PulsePal | Sanwork Inc. |
| Blue Laser SSL-473-0100-10TM-D-LED | Sanctity Laser |
| Custom-built recording microdrive for electrodes and optic fibers | Multiple companies |
| Silicon probes | Neuronexus |

### Open Ephys plug-ins

Open Ephys is an open source platform for multi-channel electrophysiology experiments (Siegle et al., 2017). Its plugin-based workflow is designed to facilitate real-time feedback in neuroscience experiments. We used the ZeroMQ interface by Francesco Battaglia (Donders Institute, Radboud University) implemented as a filter plug-in to Open Ephys, modified to support more recent Open Ephys versions. ZeroMQ is a lightweight network library that simplifies setting up some typical network topologies. The plugin broadcasts recorded data and events that can be subscribed to by external applications through ZeroMQ sockets created by ZmqInterface.cpp. The plugin uses a heartbeat mechanism to track which applications are currently connected to the data stream. The data content is dependent on the position of the ZMQ plug-in in the signal chain. We suggest sending filtered data appropriate for spike detection using the plug-in. For instance, band-pass filtering between 600-6000 Hz enables threshold-based action potential detection. Our Python interface implements thresholding itself, therefore a Spike Detector plug-in should not be included before the ZMQ interface. We use the following signal chain: Rhythm FPGA – Common Average Reference – Bandpass filter – ZMQ Interface – LFP viewer (Figure 1).

**Figure 1:**
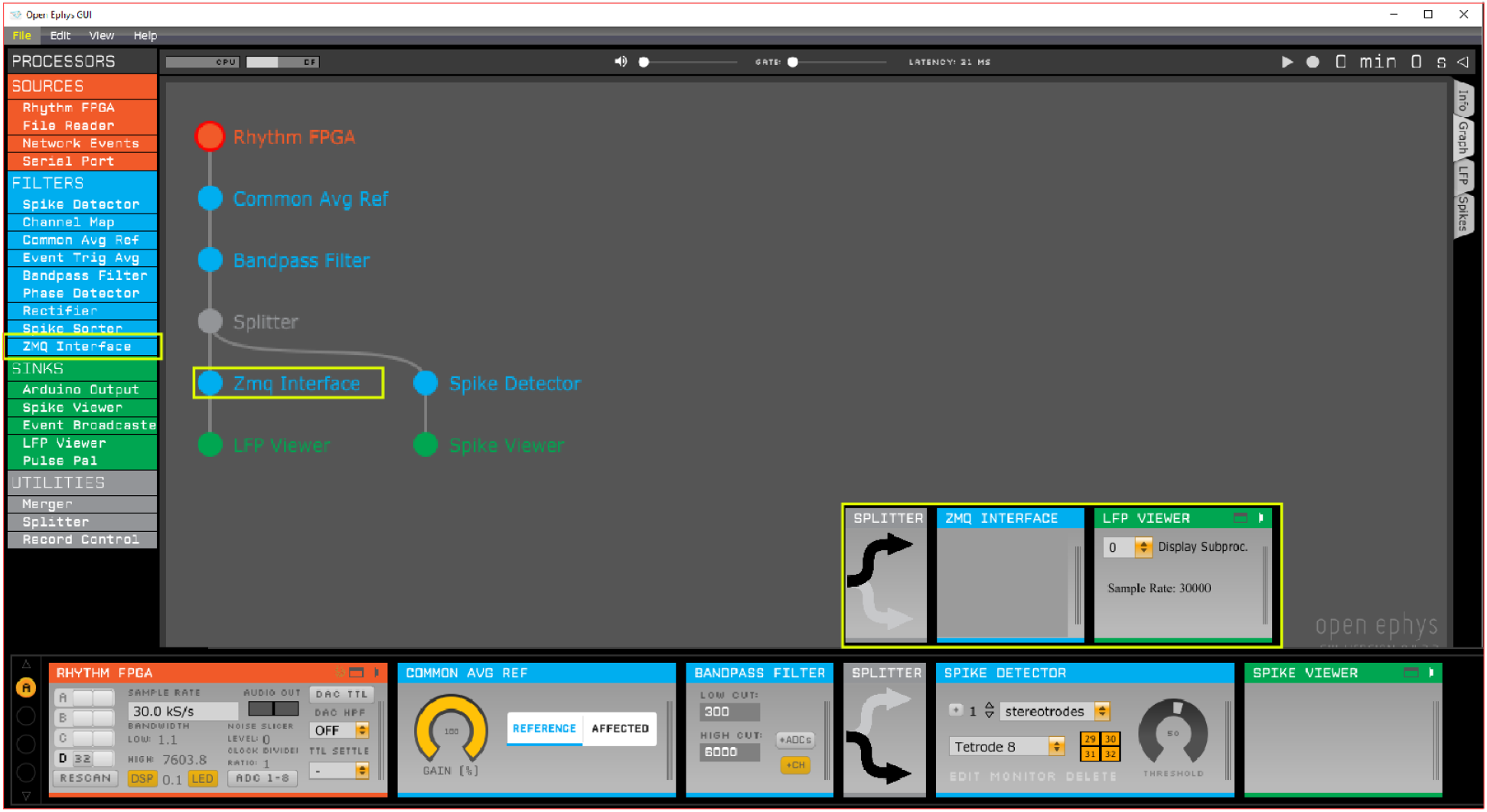
Open Ephys signal chain used to produce the figures of this work. The ZMQ interface (yellow box) is positioned after the ‘Bandpass filter’ and before the ‘LFP viewer’.

The ZMQ Interface plugin opens a ZMQ publisher socket to allow one or more ZMQ clients to subscribe (connect) locally or over the network. Though the system is typically used with a single client connected locally, it is possible to use multiple clients on multiple PCs analyzing the same Open Ephys data source simultaneously with different settings. The ZMQ plugin creates JSON format data packets from the digitized data and event metadata (e.g. timestamp, event channel, number of data channels and sample count) and sends it over to the client(s). Another socket for event messages and responses is used for heartbeat messages to inform the plugin about the connected clients.

### Python GUI

We developed a graphical user interface in Python based on pyqtgraph to visualize PETHs aligned to external events during data acquisition in real time. The GUI is compatible with Python 2.7 and Python 3 as well.

#### Histogram window

The main GUI window is handled by gui.py, which schedules data reading, spike discrimination, performs histogram calculation and enables the adjustment of parameter setup. The main window displays histograms, parameters and buttons for handling the configuration and the different plots (Figure 2).

**Figure 2:**
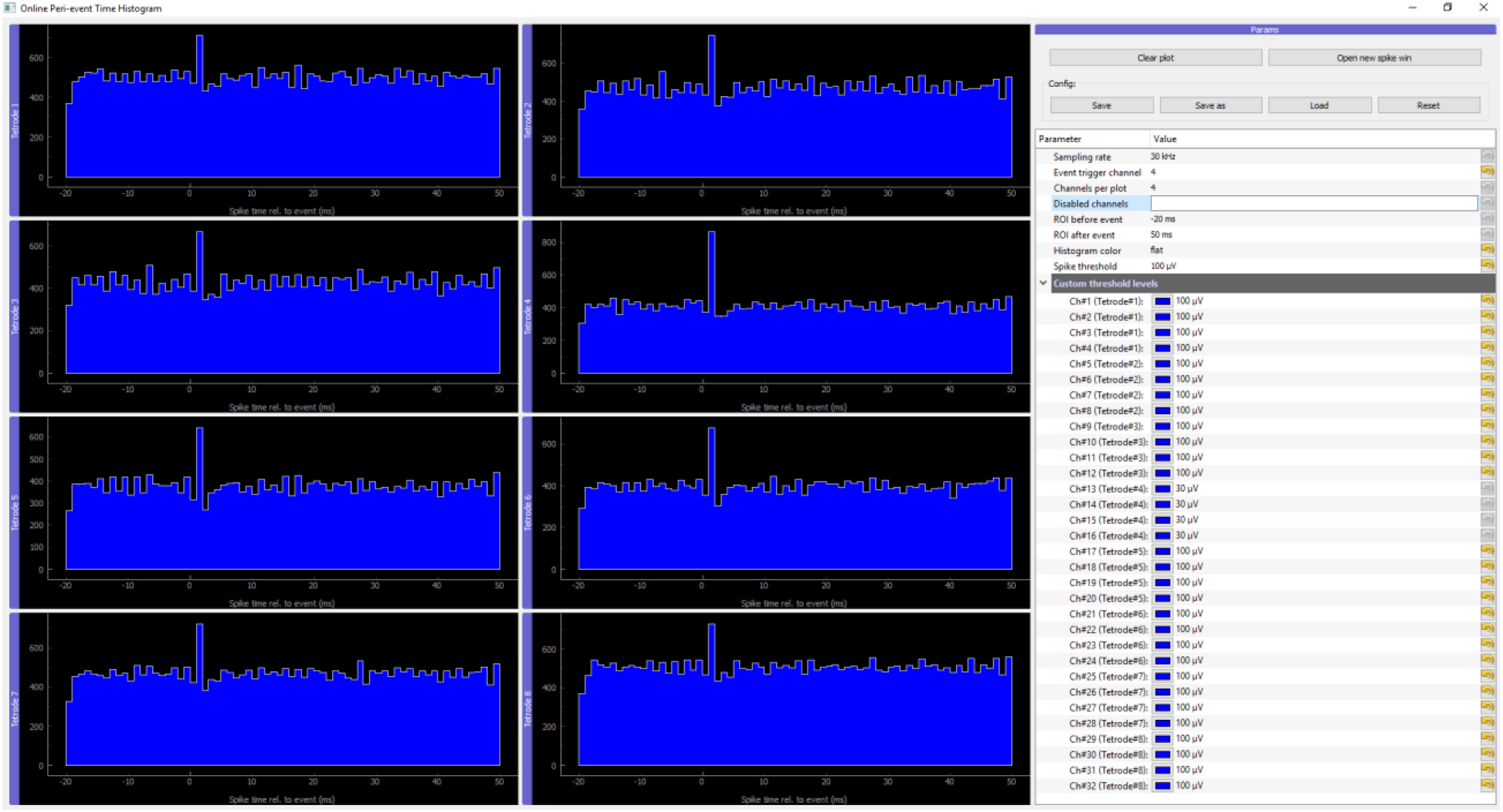
Main GUI of OPETH. The histograms of the different channels or polytrodes are displayed on the left side of the window. The menu on the right side allows changing the parameters and interacting with the GUI.

Histogram channels are collected in groups of four by default as for classical tetrode recordings, but the ‘Channels per plot’ option can be set from 1 to 8 allowing visualization for single electrodes, stereotrodes or silicone probes of different channel configurations. The different weight of each channel of the polytrode can be visualized separately in each histogram window by setting ‘Histogram color’ (Supplementary figure 1A) to ‘aggregate’ (Supplementary figure 1B). This way, the resultant graphic displays a combined multicolored histogram, showing each channel’s contribution to the histogram in a different color. If the channels of a tetrode are to be compared, it is recommended to use the ‘channels’ histogram view. In this case histograms of individual channels are not stacked, but instead overlaid in different colors with line plots (Supplementary figure 1C).

‘Sampling rate’ should be adjusted to match the samples per second of the Open Ephys ‘RHYTHM FPGA’ module. Changing the ‘channels per plot’ option automatically changes the number of histograms displayed. Briefly, ‘Clear plot’ clears all histograms and ‘Open new spike win’ initiates a new spike window (see below). The parameters can be saved and loaded for each experimental subject. The threshold options allow selection of a global threshold applied on all channels or individual thresholds for each channel. The ‘event trigger channel’ sets which Open Ephys I/O board channel is used as trigger for the histograms. The ROI options allow setting the time window before and after the trigger. Finally, ‘Disabled channels’ allows inactivating unused or noisy channels.

We implemented a way of storing all settings in configuration files in the ini file format. As it is a common use case to have multiple experimental projects running in parallel, there is a ‘Save as’ option to store the configuration in a different file. The system remembers the last stored config (file path is stored in the ‘lastini.conf’ file) and loads it automatically on startup. Config file handling is based on the configparser python module.

#### Raw analog data window

A real time data viewer window was implemented to display data received directly from Open Ephys, allowing low-level visualization of the output provided by the ZMQ plugin. Since the main purpose of this window is to provide feedback for debugging, channels are auto-scaled and thus do not provide information on actual voltage levels. The plot in the top half of the window is a one-second-long rolling display that plots all channels simultaneously (Supplementary figure 2, top). In the interest of CPU time, the plot is updated at a low frame rate and the data displayed are downsampled to 1000 Hz for this view. The bottom part of the window features a stimulus counter and presents short windows of the same analog data, uncompressed and aligned to the trigger stimuli (Supplementary figure 2, bottom). The window boundaries with respect to the trigger are set by the parameters ‘ROI before event’ and ‘ROI after event’. This window can be closed independently of the main window if not required.

#### Spike window

Spike waveforms triggered on TTL can be visualized in separate Spike windows. Spike windows can be opened from the main histogram window and are handled by spike_gui.py. Each window displays spikes of a single channel; the selected channel can be changed real time. The plot displayed in the top part of the window shows the raw input data of the channel aligned to the event, with the detected spikes overlaid in color. These detected spikes are enlarged separately in the bottom part with the same color code. The raw data display of the Spike window presents uncompressed data within the region of interest around the trigger (as set in parameters). The spike plots display a short segment of data before and after the peak value (red dot) of the spikes (−0.3 ms to +1 ms by default). If the ‘Update only on spike’ option is selected, spike windows are updated when new spikes are detected within the ROI of the trigger; otherwise, spike windows are updated 5 times per second even when no spikes are present. Multiple Spike windows can be displayed simultaneously, however, this is CPU intensive and opening too many Spike windows will slow down the application (Figure 3).

**Figure 3:**
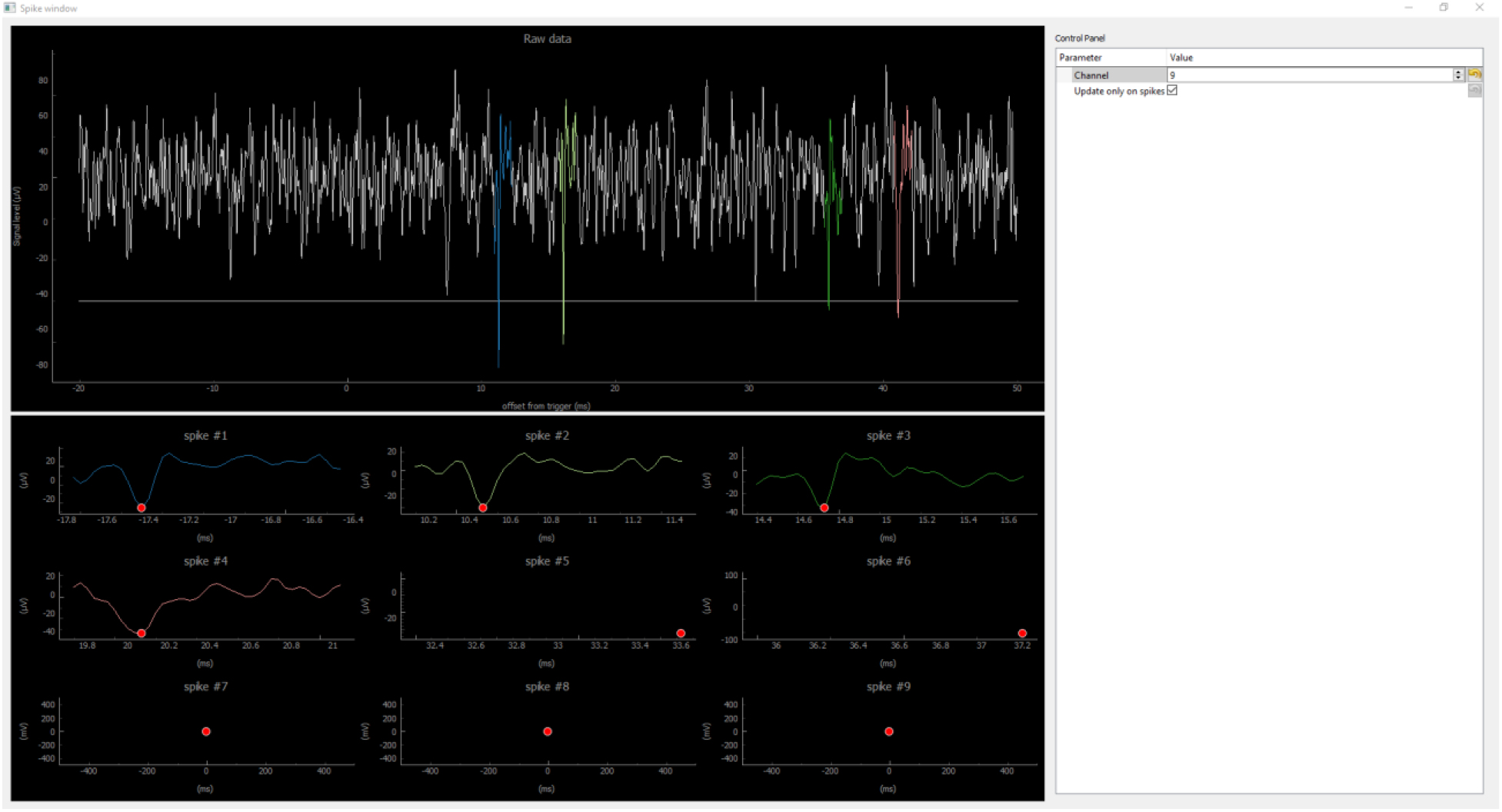
Spike window. Top, continuous data from the selected channel with spikes detected in restricted temporal windows (ROI) around the trigger TTLs, overlaid in color. Horizontal line shows the spike detection threshold. Bottom, zoomed-in windows for the same detected spikes.

### Operation overview

Until the Open Ephys ZMQ plugin connection is established, the GUI displays “Awaiting data”. Once the first chunk of data is received, the exact GUI layout is determined based on the number of channels and the histogram plots are displayed.

Input data arriving from Open Ephys is handled by comm.py, which takes care of parsing the JSON structures containing the measurement samples and trigger events. Depending on the type of the parsed input data, trigger events are stored in OpenEphysEvent objects (defined in openephys.py) and sample data are stored directly in a 2D circular (or rolling) buffer implemented in circbuff.py; the data flow is managed by the Collector class in colldata.py (see Figure 4). The openephys.py and some of the comm.py interface routines are based on the python samples created by Francesco Battaglia, while we have developed the circular buffer and colldata.py data handling methods from scratch as well as the entire visualization UI.

**Figure 4:**
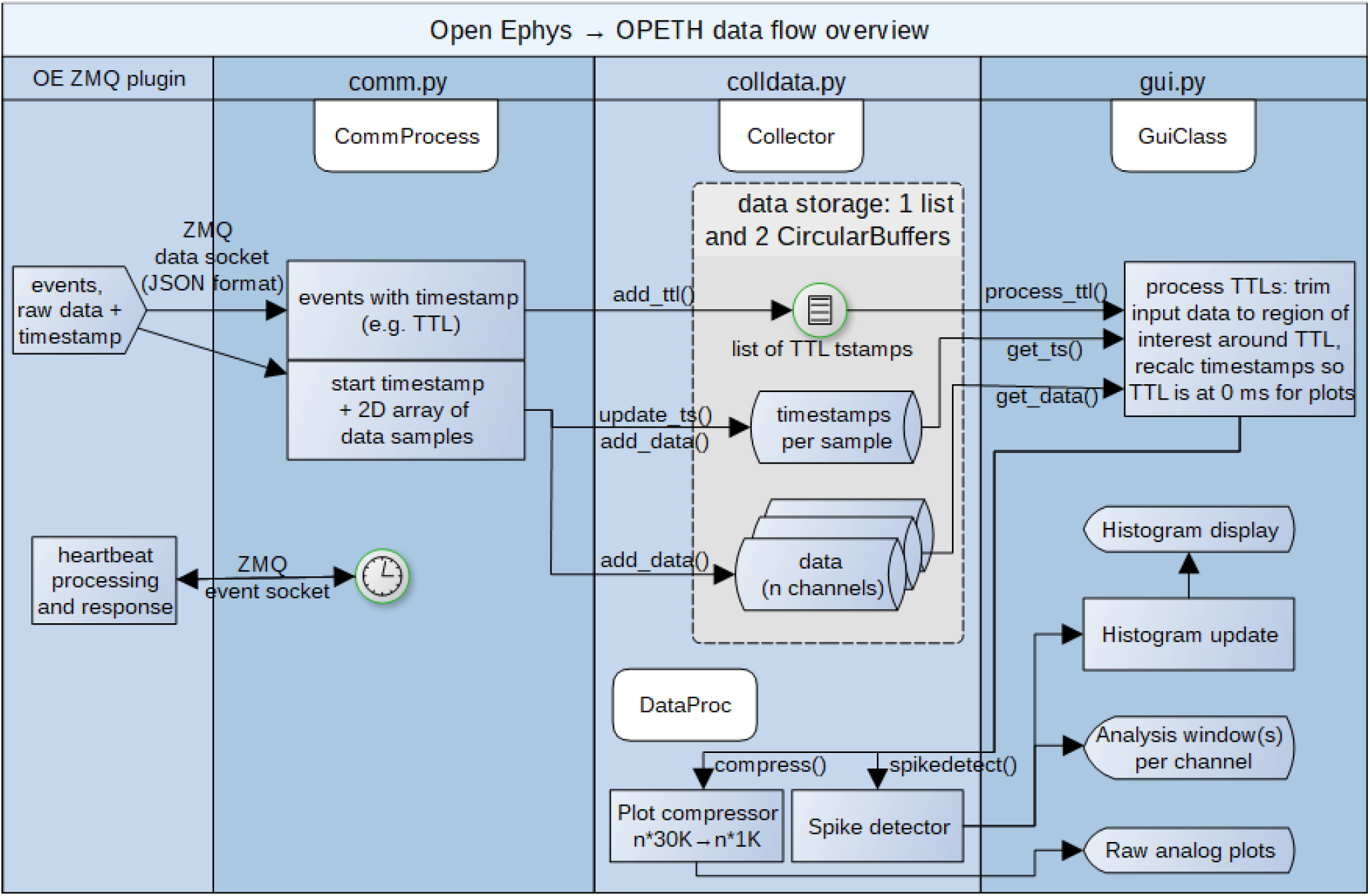
Simplified diagram of data flow.

The circular buffer allows storing the last few seconds of data for the raw data displays, which gets downsampled by the compress method of DataProc class in colldata.py to save CPU time when plotting all signals simultaneously. Only minimum and maximum values are displayed for every 30 data points. The compressed data is used in the raw analog data window only. The spike detection is performed on the original data as discussed in detail later.

The data collection, spike detection and all plot updates are scheduled by the main loop in gui.py by the update() method that is periodically called by the Qt environment. We note that while the current implementation is single threaded except for ZMQ messaging, it is worth exploring options for multiprocessing in future updates. Figure 4 shows a schematic version of the data flow.

### Spike detection

We created a spike discrimination routine independent of Open Ephys spike detectors that resides in the DataProc class of colldata.py. The spikedetect() is called by the update() routine whenever a new TTL signal or other event is detected on the currently selected trigger channel. It performs simple spike discrimination based on voltage levels within the region of interest (ROI) around the stimulus or timestamped event. Whenever a spike is detected because of exceeding the threshold level, new spikes are not detected until a predefined holdoff time is passed and the voltage level drops below threshold again. The implementation works by default with negative threshold levels to allow extracellular spike detection on non-inverted raw voltage data. (This can be modified via the `NEGATIVE_THRESHOLD` constant of gui.py). As higher level spike source identification was not a target of our tool, spikes are identified per channel and are not sorted or verified across multiple channels. Overlapping trigger ROIs may result in repeatedly detected spikes in the overlapped region (i.e. some spikes appear twice in the histogram).

Spike detection is done in two phases: first the input data is thresholded by comparing the analog input levels to the threshold parameter, then the thresholded array is processed channel by channel. For each spike on the given channel the starting and final position exceeding the threshold is determined, and between them the maximum or minimum value of the input data (depending on spike polarity) gives the spike position.

If the spike detection happens on negative threshold, the falling edge of the input signal is to be detected, in which case the positive threshold value of the GUI setting is automatically inverted for the spikedetect function. The thresholding is performed in a single step for all channels, with possibly unique threshold levels per channel. This results in a 2D array with one row per channel, each column containing Boolean values indicating whether the given sample is exceeding the rising/falling edge threshold level. Non-disabled channels are searched one by one for spikes in a loop. The code excerpts below show the negative threshold case with local minimum search as depicted on Figure 5.

**Figure 5:**
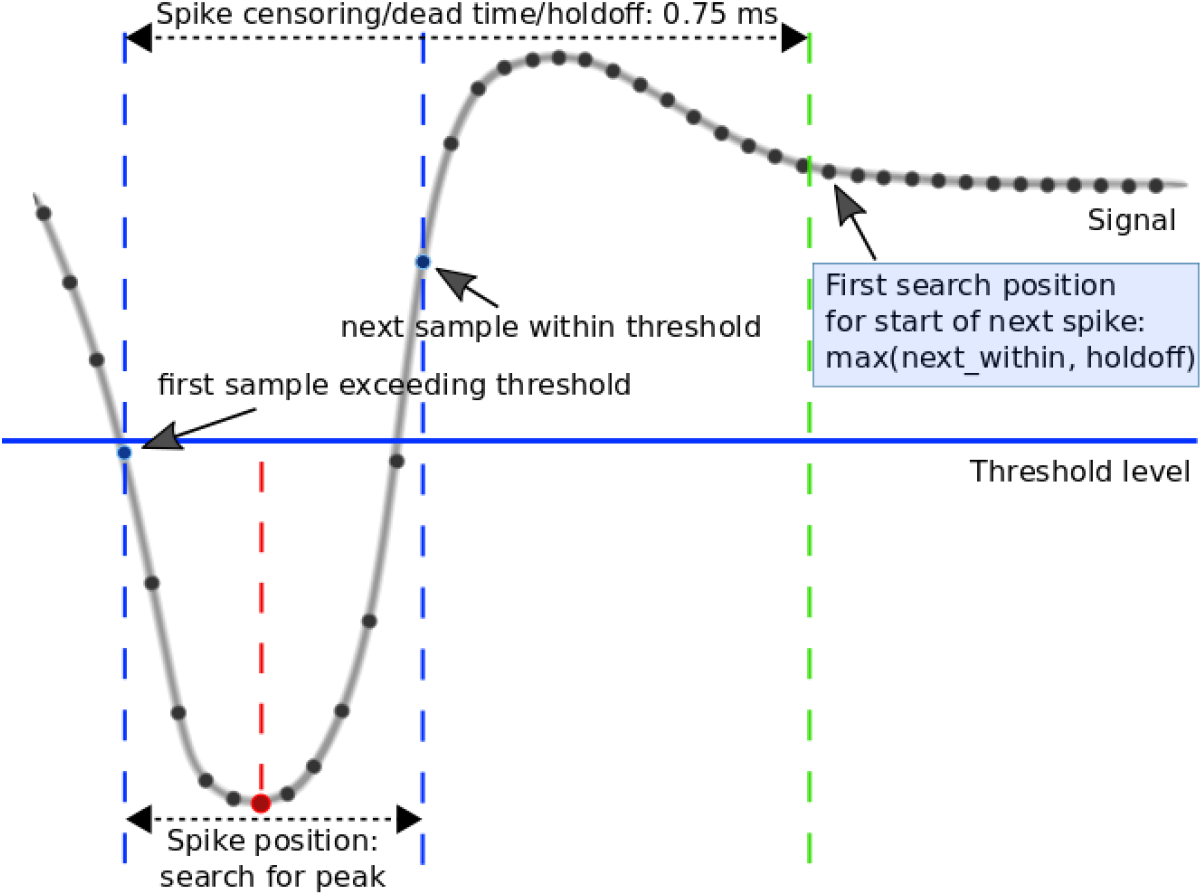
Schematics of spike detection. Key variables for spike detection based on threshold crossings are indicated. Note that sample count does not correspond to default 30kS/s.

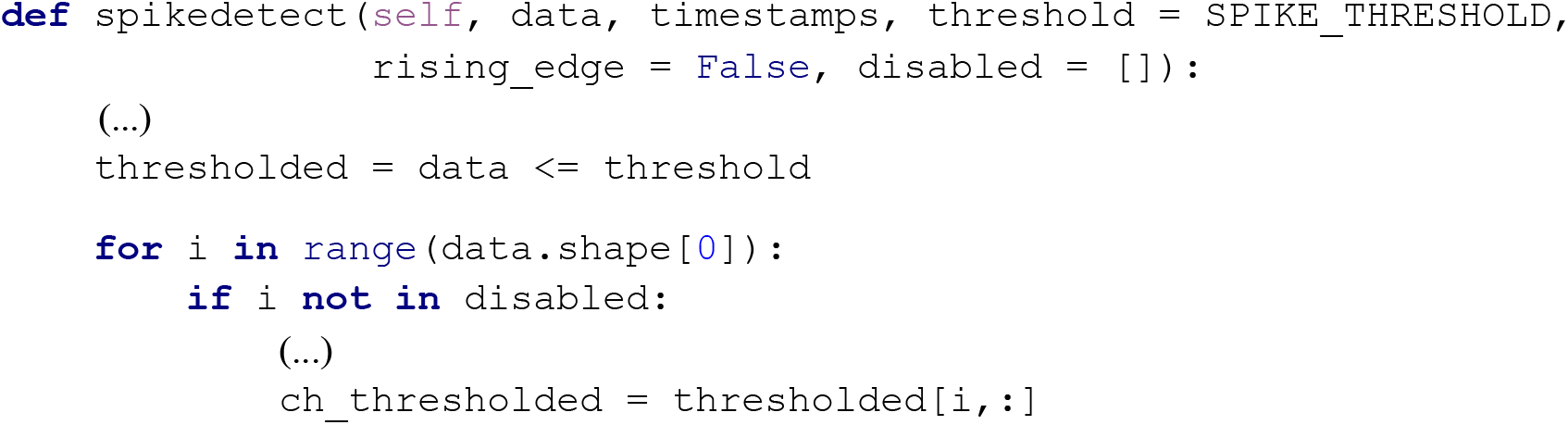

To detect a spike we look for the first False→True transition in the thresholded array for the starting position (`first_over_thresh`). The end of the spike is the next nearest True → False transition in the thresholded data (`next_below_thresh`). Search for the next spike starts from this end position (`offset`), following a predefined dead time/censoring period of 0.75 ms by default (`SPIKE_HOLDOFF_SAMPLES`) to avoid repeated detection of the same spikes. The spike position (position of the negative peak or minimum value) and timestamps are then collected into lists for the spikedetect function return values with one list of spike positions per channel.

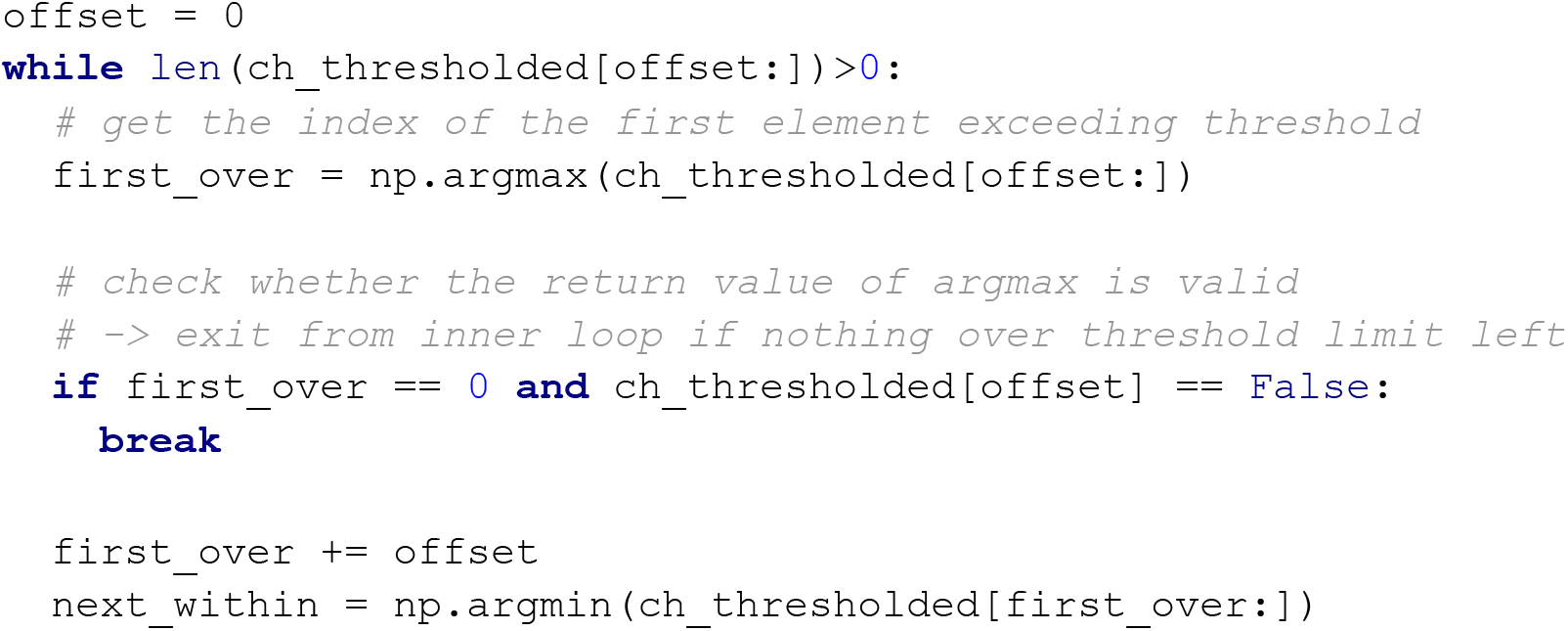
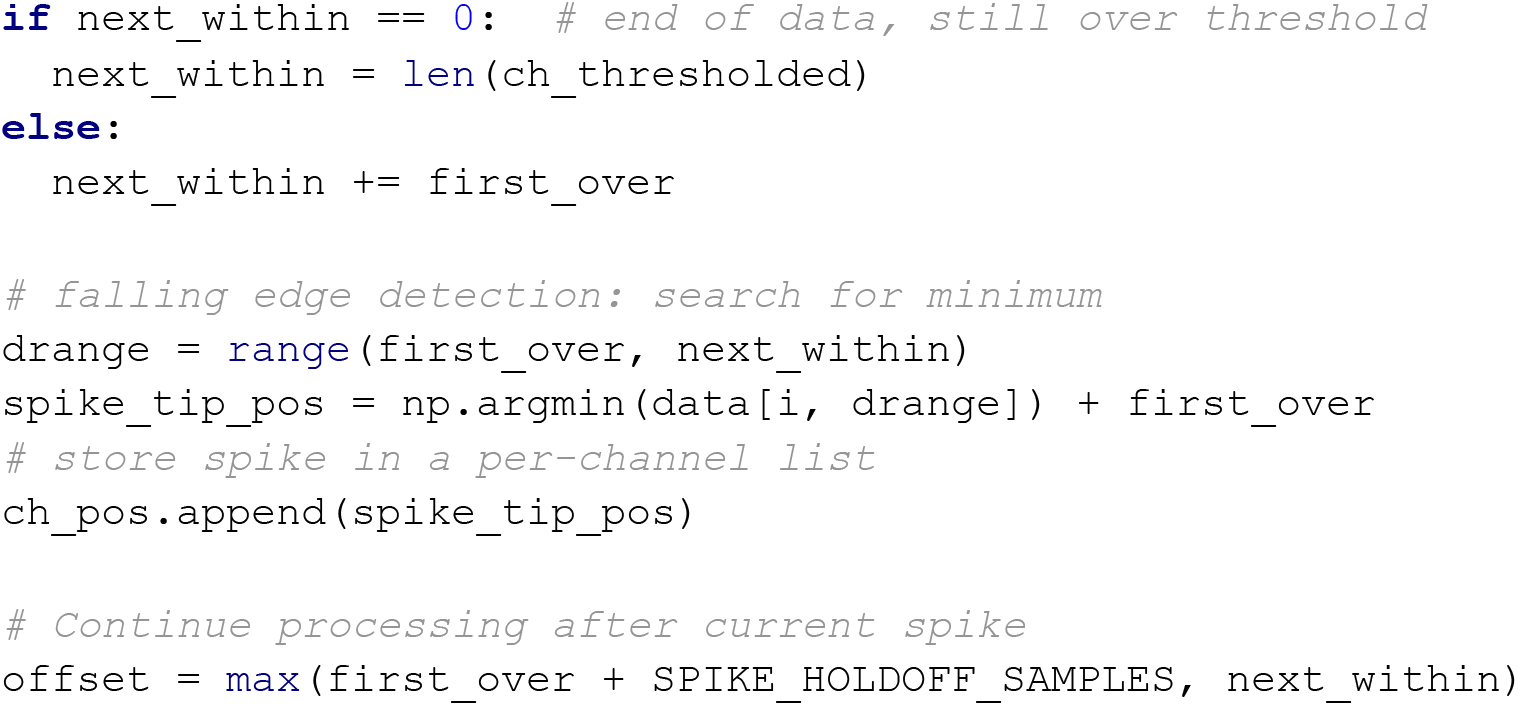

Spike thresholds can be adjusted individually for each channel or for all channels at the same time, using either µV or mV as unit of measurement. The region of interest around events for spike detection is set in milliseconds. It is possible to disable channels or sets of channels from spike detection, e.g. in case of broken channels. Such channels can be listed either comma-separated or with the dash notation (1, 2, 3 or 1-3) in the disabled channels input field. The histograms of the disabled channels are not updated.

### In vivo experimental procedures

#### Animals

Electrophysiological, optogenetic and behavioral data showed in this study were obtained from two adult female mice (BAC-Vglut2-IRES-Cre, C57Bl/6J and ChAT-IRES-Cre, B6129F1). All experiments were approved by the Committee for the Scientific Ethics of Animal Research of the National Food Chain Safety Office and were performed according to the guidelines of the institutional ethical code and the Hungarian Act of Animal Care and Experimentation (1998; XXVIII, section 243/1998, renewed in 40/2013) in accordance with the European Directive 86/609/CEE and modified according to the Directives 2010/63/EU.

#### Surgery and virus injection

Mice were anesthetized with an intraperitoneal injection of ketamine-xylazine (0.166 and 0.006 mg/kg, respectively). The scalp was shaved and disinfected (Betadine) and local anesthetics was applied subcutaneously (Lidocaine). The mouse was positioned in the stereotaxic frame and the eyes were protected with eye ointment (Laboratories Thea). The skin was removed above the calvaria and the skull was cleaned; the head of the animal was leveled using Bregma and Lambda (Paxinos et al., 2001) and lateral points equidistant from the sagittal suture.

In the Vglut2-Cre animal, a cranial window was opened in order to access the medial septum (MS) with a 10º lateral angle (MS 10º, antero-posterior +0.90 mm, lateral, 0.90 mm). An adeno-associated virus vector allowing Cre-dependent expression of channelrhodopsin2 [AAV 2/5. EF1a.Dio.hChR2(H134R)-eYFP.WPRE.hGH] was injected into the MS at 3.95, 4.45 and 5.25 mm depth from skull surface (200 nl at each depth). The skin was sutured; local antibiotics (Neomycin) and a subcutaneous injection of analgesic (Buprenorphine 0.1 mg kg^−1^) were applied.

In the ChAT-Cre animal a craniotomy was performed above the horizontal nucleus of the diagonal band of Broca of the basal forebrain (HDB, antero-posterior 0.75 mm, lateral 0.60 mm) and the same virus was injected into the HDB at 5.00 and 4.70 mm depth from skull surface (300 nl at each depth). Additional holes were drilled above the parietal cortex for ground and reference. The surface of the skull was covered with a thin layer of Super-Bond C&B (Sun Medical) and a custom-built microdrive (Hangya et al., 2015; Kvitsiani et al., 2013) with 8 tetrodes was implanted in the targeted area. The microdrive-skull junction was protected with Kwik-Cast sealant (World Precision Instruments). The microdrive was secured to the skull with dental acrylic resin (Lang Dental). A titanium headbar was also attached to the skull to allow headfixation. Analgesic and antibiotics were applied as above.

Mice were allowed to recover for ten days, receiving subcutaneous injections of analgesic (Buprenorphine 0.1 mg kg^−1^) and local application of antibiotics (Neomycin) as necessary.

#### Anesthetized recordings

Twenty days after the virus injection the Vglut2-Cre animal was anesthetized with an i.p. injection of 20% urethane (Sigma-Aldritch, 0.007 ml g^−1^ body weight). The depth of anesthesia was evaluated by pinching the paw or ear of the animal. When there were no reflexes elicited by the pinching, the throat was shaved and topical lidocaine was applied. A tracheotomy was performed in order to sustain a constant airflow (Moldestad et al., 2009). The animal was placed in a stereotaxic frame and, after opening the skin and leveling the skull, trephine holes were made above the MS (silicon probe MS 10º, antero-posterior, +0.90 mm, lateral, 0.90 mm; optic fiber MS 5º contralateral, antero-posterior, +0.90 mm, lateral, −0.50 mm), the hippocampus (silicon probe HPC, antero-posterior, −2.20 mm, lateral, 1.50 mm) and two above the cerebellum for reference electrodes. A Neuronexus A1×32-6mm-50-177-CM32 silicon probe was placed in the hippocampus at 2.20 mm depth from skull surface, and a Neuronexus Buzsaki32-H32_21mm probe was lowered to the dorsal boundary of the MS at a 10º lateral angle (3.95 mm from skull surface). Reference electrodes for both probes were placed in the cerebellum and ground electrode was placed in the spinotrapezius muscle. A 200 µm core optic fiber was lowered 500 µm above the shanks of the MS probe. The MS probe and the optic fiber were lowered in 100 µm steps for recording, spanning the entire depth of the MS. Extracellular data were collected by the Open Ephys data acquisition system, digitized at 30 kS/s. Each recording session consisted of an optical tagging period of two minutes, followed by a baseline period of five minutes. Three consecutive repetitions of one-minute tail pinch induced theta activity followed by one-minute control recording were applied, finishing the recording session with another two minutes length optical tagging period. After each recording session, the MS probe and optic fiber were lowered 100 µm.

#### Head-fixed recordings and behavioral procedures

Once fully recovered, the drive-implanted Chat-Cre animal was trained on a head-fixed auditory cued outcome task implemented in a go/no-go paradigm. Briefly, the animal was water restricted for three days. On the fourth day, the animal was head-fixed in the behavioral environment (Solari et al., 2018), where after a few free water delivery trials, a go tone (10 kHz, 50 dB, 1 s) was presented. Licking during the tone resulted in the release of a 3 µl water droplet as reward. Once the animal was familiarized with this paradigm, a second tone (4 kHz, 50 dB, 1 s) was introduced, predicting the delivery of an air puff (duration, 200 ms). In the final task, a balanced mixture of the two tones were randomly interleaved in which the 10 kHz tone predicted expected reward in 80% of trials, unexpected punishment in the 10% of the trials and omission in the remaining 10% of trials; the 4 kHz tone predicted expected punishment the 65% of trials, unexpected reward in the 25% of the trials and omissions in the remaining 10%. Extracellular data were collected during task performance by the Open Ephys data acquisition system, digitized at 30 kS/s.

#### Data analysis

Offline data analysis was performed using built-in and custom-built Matlab (Mathworks) scripts. Action potentials were manually sorted into putative neuronal clusters based on amplitude (peak-to-valley), waveform energy and first principal component features using the MClust software (A. D. Redish). L-ratio (<0.05) and isolation distance (>20) were taken as cluster quality measures (Schmitzer-Torbert et al., 2005).

Vglut2 positive neurons were identified by optogenetic tagging (Kvitsiani et al., 2013; Lima et al., 2009; Pi et al., 2013). Significant light activation was assessed by the Stimulus-Associated spike Latency Test (SALT), using the code available at http://kepecslab.cshl.edu/salt.m (Kvitsiani et al., 2013).

## Results

In the following, we provide two example applications of OPETH. First, we used OPETH for online optogenetic tagging of medial septal glutamatergic neurons in an acute anesthetized experiment. Second, we performed chronic recordings from the horizontal nucleus of the diagonal band of Broca (HDB) while a mouse performed a head-fixed auditory cued outcome task. We demonstrated the presence of punishment-activated HDB neurons during the recording online with OPETH, and later confirmed this result by offline analyses.

### Real-time optogenetic tagging

Optogenetic tagging allows the identification of neuron types in extracellular recordings performed in transgenic animals. For instance, we use optogenetic tagging in acute anesthetized experiments to investigate the role of different genetically defined types of medial septal (MS) neurons in the genesis of neural oscillations and network synchrony (Buzsáki and Moser, 2013; Hangya et al., 2009; Wang, 2002). However, the yield of such experiments can greatly be increased if the presence of optogenetically tagged neurons can be established online during the recording. Additionally, real-time feedback helps reducing artifacts in the recordings introduced by photostimulation.

Therefore, we tested OPETH in this *in vivo* optogenetic tagging experiment. A BAC-Vglut2-IRES-Cre mouse previously injected with a viral construct allowing Cre-dependent expression of the light sensitive channelrhodopsin2 protein in glutamatergic MS neurons was anesthetized with urethane. A 32-channel linear silicone probe was placed in the hippocampus for local field potential recordings and a 32-channel four-shank silicon probe was lowered into the MS for extracellular recording of MS units. In addition, an optic fiber was placed in the MS above the recording probe to deliver laser light for photostimulation. Throughout the experiment, laser-triggered responses of the MS neurons were monitored by the OPETH *Histogram Window* (Fig. 6A). Once putative light-evoked action potentials were detected, the *Channel view* was used to assess the channel with the largest light effect (Fig. 6B). *Spike Window* for this channel was then opened and monitored while adjusting light intensity levels to avoid photoelectric effects (Fig. 6C-D). This protocol allowed us to ‘hunt’ for optogenetically identified glutamatergic MS neurons with removing the potential confounds arising from photostimulation-related electrical artifacts, increasing the efficiency of the experiment.

**Figure 6:**
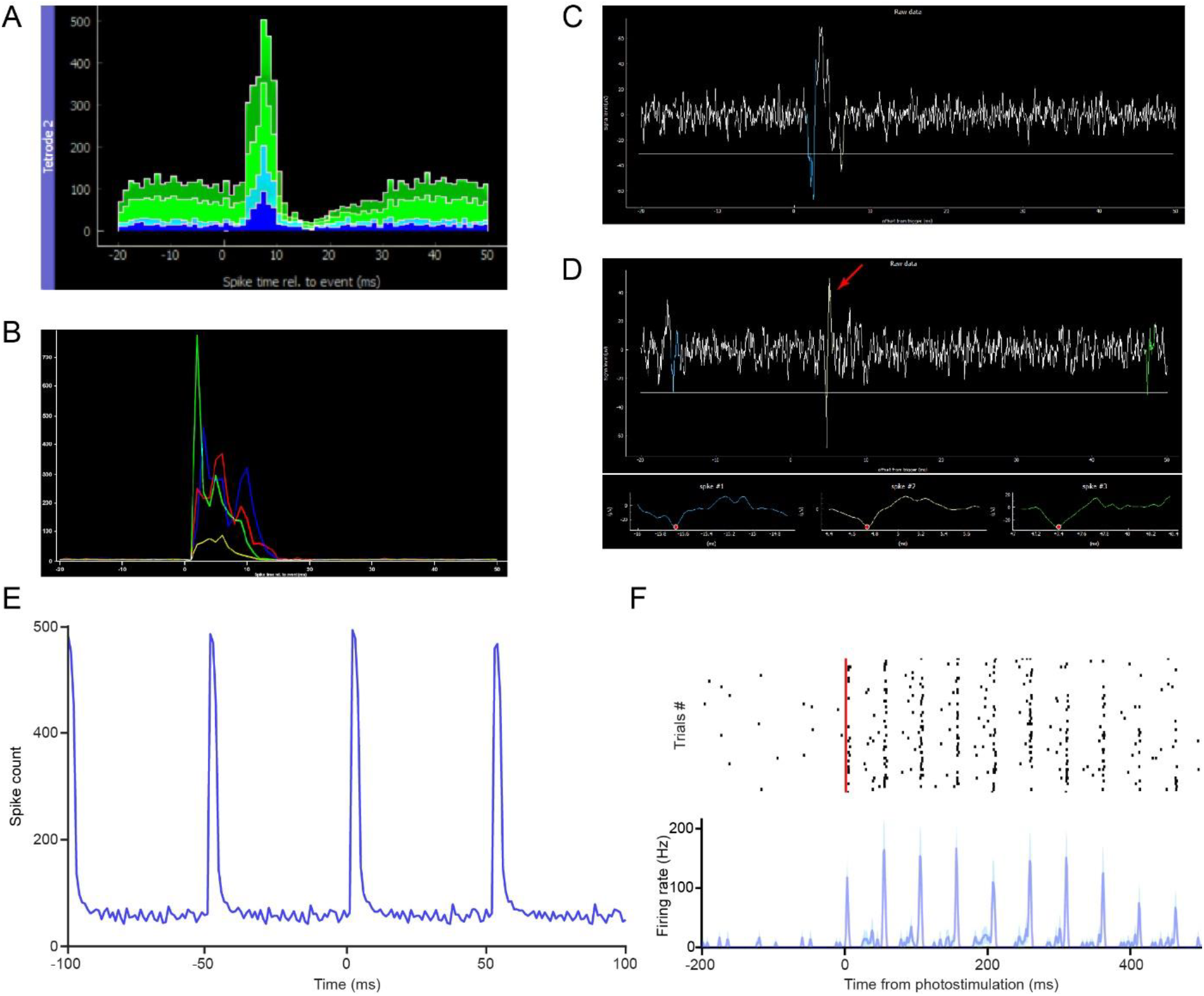
*In vivo* optogenetic tagging experiment for testing OPETH. A) *Histogram Window, ‘aggregate view’* showing neural responses to laser pulses in a Vglut2-Cre mouse expressing channelrhodopsin2. B) *‘Channel view’* was used to determine the most responsive channel of the tetrode of interest. C) *‘Spike window’* of the selected channel showing a photoelectric artifact evoked by high intensity laser stimulation. D) The *Spike Window* allowed us to tune down the light intensity until the putative tagged cell was not masked by artifacts (red arrow). E) The offline peri-event time histogram showed strong light-evoked activity on the selected tetrode. F) Spike raster and peri-event time histogram of a responsive MS neuron aligned to the onset of the laser pulse train, from the channel selected via OPETH.

After concluding the experiment, we performed offline peri-event time histogram analysis, aligned to the onset of the photostimulation pulses. This confirmed the presence of light responses on the same tetrodes as shown by OPETH (Fig. 6E). Finally, spike sorting of three recording sessions revealed significantly light-activated neurons recorded by the same tetrodes as indicated by OPETH (n = 10, p < 0.001, Fig. 6F).

### Real-time peri-event time histogram in behaving mice

In addition to optogenetic tagging, OPETH enables online tracking of neural responses to behaviorally relevant external events such as cue stimuli and reinforcement. To demonstrate this, we next tested OPETH’s ability to detect neuronal activity changes during a head-fixed go/no-go task in an awake behaving mouse.

A mouse was fully trained on a head-fixed auditory cued outcome task, in which two pure tones of different pitch signaled different outcome probabilities, predicting either likely reward (water) vs. surprising punishment (a puff of air) or vice versa. Two different TTL pulses were sent to the Open Ephys I/O board every time reward or punishment was delivered, allowing OPETH to visualize the neuronal response to each of the behavioral outcomes. The mouse performed a total of 254 trials in a single recording session. Throughout this session, the *Histogram Window* of OPETH clearly showed a neuronal response to punishment on most tetrodes (Fig. 7A), while no response to reward delivery could be detected (Fig. 7B). We noted that this punishment response was already detectable after the first few punished trials, showing the sensitivity of detection by OPETH.

**Figure 7:**
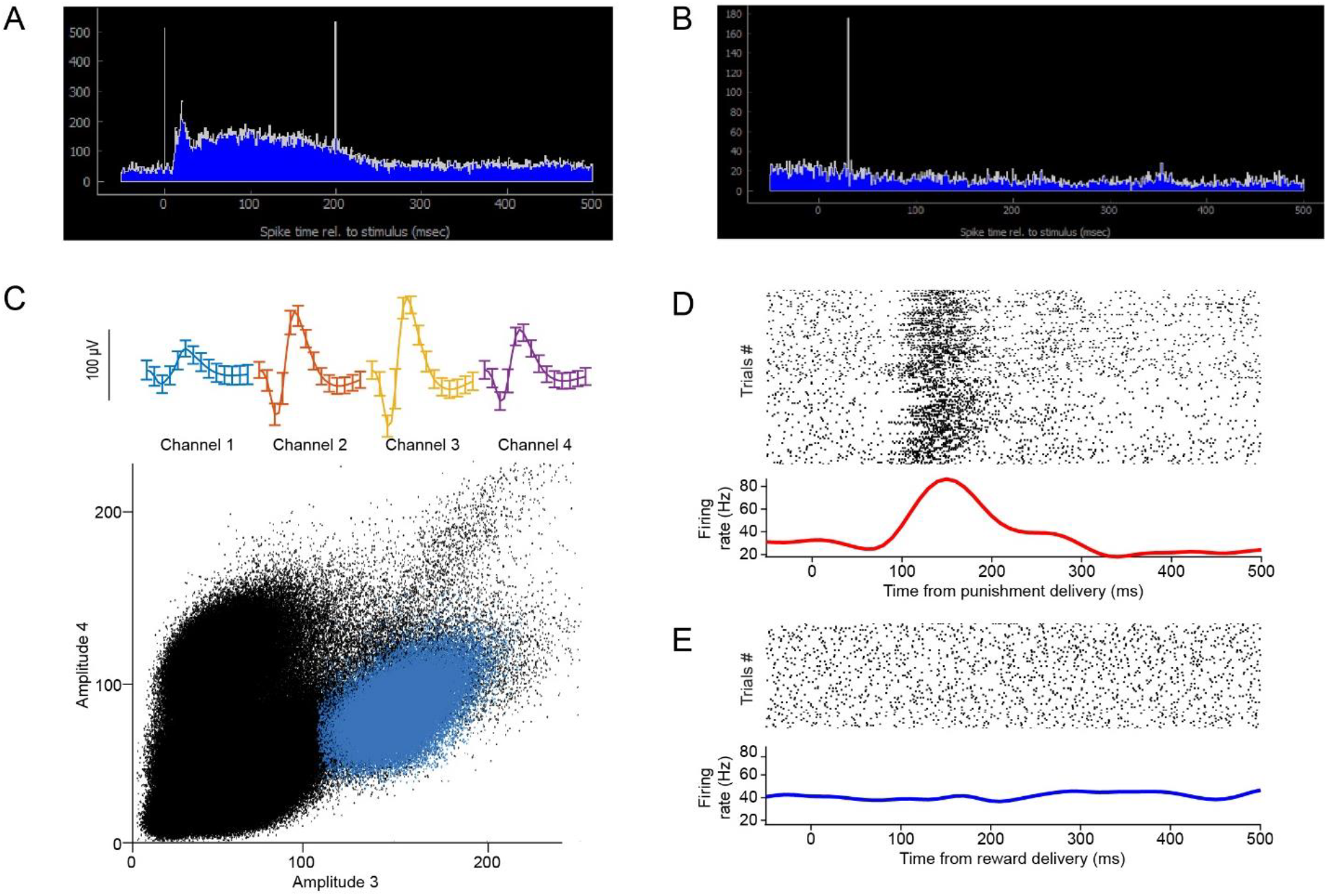
Neuronal responses to behaviorally relevant events detected real-time by OPETH. A) Most of the tetrodes showed an increase in neuronal activity in response to punishment delivery (vertical lines indicate artifacts due to valve opening and closing). B) No neuronal response was detected after reward delivery. C) Spike shape and cluster projection of an example punishment-activated reward-unresponsive neuron after offline spike sorting. D) Spike raster and PETH aligned to the onset of punishment delivery. E) Spike raster and PETH aligned to the onset of reward delivery.

After the recording session, offline analysis confirmed the presence of neurons that responded selectively to punishment, as expected based on the online feedback by OPETH. Fig. 7C-E shows a well isolated neuron that responded with an increase of firing after air puff punishment, but not after water reward.

## Discussion

Real-time feedback while performing electrophysiology recordings is important to guide decisions during the experiment. Here we described OPETH, an open source online tool for providing such feedback by visualizing peri-event time histograms. In addition, we demonstrated its usefulness when conducting optogenetic tagging or behavioral experiments combined with single cell or multiunit recording. OPETH is based on Open Ephys, an open source data acquisition system including software (Siegle et al., 2017).

### Open source

There is an increasing number of open source tools in neuroscience, which is also paralleled by an increased awareness of the open source movement in general (Gleeson et al., 2017). An important example is Open Ephys, enabled by the development of Intan chips that allowed an affordable upscaling of electrophysiology experiments. Combined with open source tools for behavior control (Sanders and Kepecs, 2012), stimulation (Sanders and Kepecs, 2014) and full behavioral environments (Devarakonda et al., 2016; Erlich et al., 2011; Solari et al., 2018), this array of recent tools has changed the way electrophysiology experiments are performed. We provide OPETH as a new member of this family that parallels the richness of features of commercial solutions (e.g. Neuralynx Histogram Display, https://neuralynx.com/software/category/sw-acquisition-control), at the same time available to the entire neuroscience community.

### Application: real-time in vivo cell type identification

In neuroscience, many of the key insights were gained by recording the electrical activity of neurons (Sviatkó and Hangya, 2017). An instructive example was the mapping of basal ganglia neurons while monkeys were engaged in a variety of behavioral tasks (DeLong, 1971; Delong et al., 1984). DeLong and colleagues performed basic linear convolution-based data analysis in the form of raster plots and peri-event time histograms, which still remains the mainstay of systems neuroscience. Eventually, these results lead to the Deep Brain Stimulation surgeries during which stimulating electrodes are lowered to the subthalamic nucleus of the basal ganglia in Parkinson’s patients, largely alleviating their otherwise often crippling motor impairments. However, the lack of proper tools to identify the great diversity of anatomically, histochemically and hodologically defined cell types of the basal ganglia *in vivo* stalled further progress (Sviatkó and Hangya, 2017).

This was first overcome by glass pipettes that allowed filling of the recorded cells by applying current pulses, called juxtacellular recording (Pinault, 1996). Then, the recent advent of imaging and optogenetic techniques (Ghosh et al., 2011; Park et al., 2015; Shin et al., 2017) opened the way to high-throughput cell type identification in awake, behaving rodents (Al-Hasani et al., 2015; Miller et al., 2019; Wang et al., 2019). This necessitates the development of new software tools aligned to this task, enabling significant increases of experimentation efficiency. OPETH provides a way of online tracking cellular responses to light flashes, in order to optogenetically identify those neurons that respond by short latency. This allows determining whether the target area has been reached, and good quality recordings of identified units can be performed. Therefore, by enabling ‘hunting’ for neurons of interest, this tool can efficiently increase the yield of optogenetic tagging experiments.

### Application: online tracking of response properties to behaviorally relevant events

Peri-event time histograms usually represent the first-pass analysis of neuronal activity of behaving animals (Endres et al., 2008; Shimazaki and Shinomoto, 2010). We have demonstrated here that this first-level analysis can be performed online, providing immediate feedback on the responsiveness of the recorded population. This may be especially useful when looking for neurons with a particular response profile, or cell types that can be identified by features of their responses. Since areas may differ significantly in the proportion of neurons responding to different sensory cues, OPETH may also allow the rough identification of target areas. Other applications include online receptive field mapping (Froemke et al., 2007) or precise localization along the frequency axis of auditory cortical tonotopy maps (Hromádka et al., 2008).

### Conclusions and future directions

By providing online access to event-aligned linear data statistics, OPETH also opens the door to more advanced online analysis. For instance, dopaminergic neurons in VTA may be identified by principal component analysis of their PETH aligned to reward and reward-predicting cues, as demonstrated by Cohen and colleagues (Cohen et al., 2012) and later applied by other labs (Takahashi et al., 2016). Therefore, adding this analysis to OPETH may allow online identification of dopaminergic cells without performing optogenetics. Other examples include online analysis of delay activity in working memory tasks, or correlating neuronal firing with reward expectations or prediction errors.

OPETH also allows future implementation of closed-loop protocols. Closed-loop approaches are gaining momentum as part of experimental procedures (El Hady, 2016) as well as in clinical applications (Ghasemi et al., 2018). Closed-loop neuronal recording in behavioral tasks has been used for assessing the role of the mouse primary visual cortex during navigation (Saleem et al., 2013), enhancing spatial navigation skills of mice by optical manipulation of the hippocampal theta oscillation cycles (Siegle and Wilson, 2014), determining the causal involvement of sharp wave ripple events in learning (Rangel Guerrero et al., 2018) and to control Drosophila feeding behavior (Moreira et al., 2019). OPETH can be used as a programmable open-source tool for closed-loop paradigms based on the detected neuronal activity, allowing high-precision automatic control of the desired output.

## Acknowledgements

We thank Nicola Solari for insightful comments on the manuscript. This work was supported by the ‘Lendület’ Program of the Hungarian Academy of Sciences (LP2015-2/2015), NKFIH KH125294 and the European Research Council Starting Grant no. 715043 to BH. BH is a member of the FENS-Kavli Network of Excellence.

## Author contributions

BH conceived the project. AS developed the OPETH software, SMB and PH performed the experiments and data analysis, SMB and AS prepared the figures, BH, AS and SMB wrote the manuscript with inputs from PH.

---

**Supplementary figure 1:**
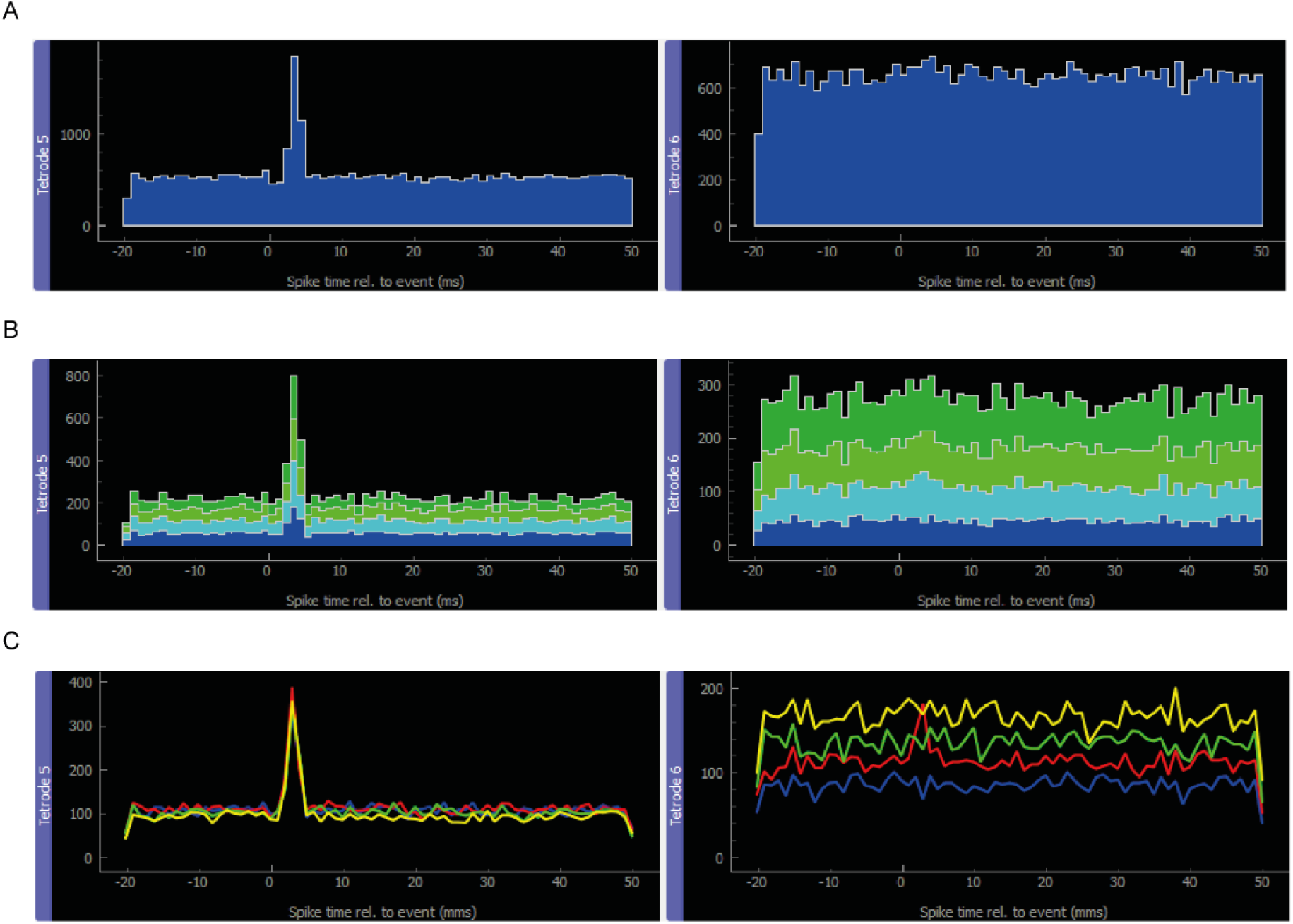
Different visualization modes of the main GUI window. A) Histogram color, B) Aggregate and C) Channels views.

**Supplementary figure 2:**
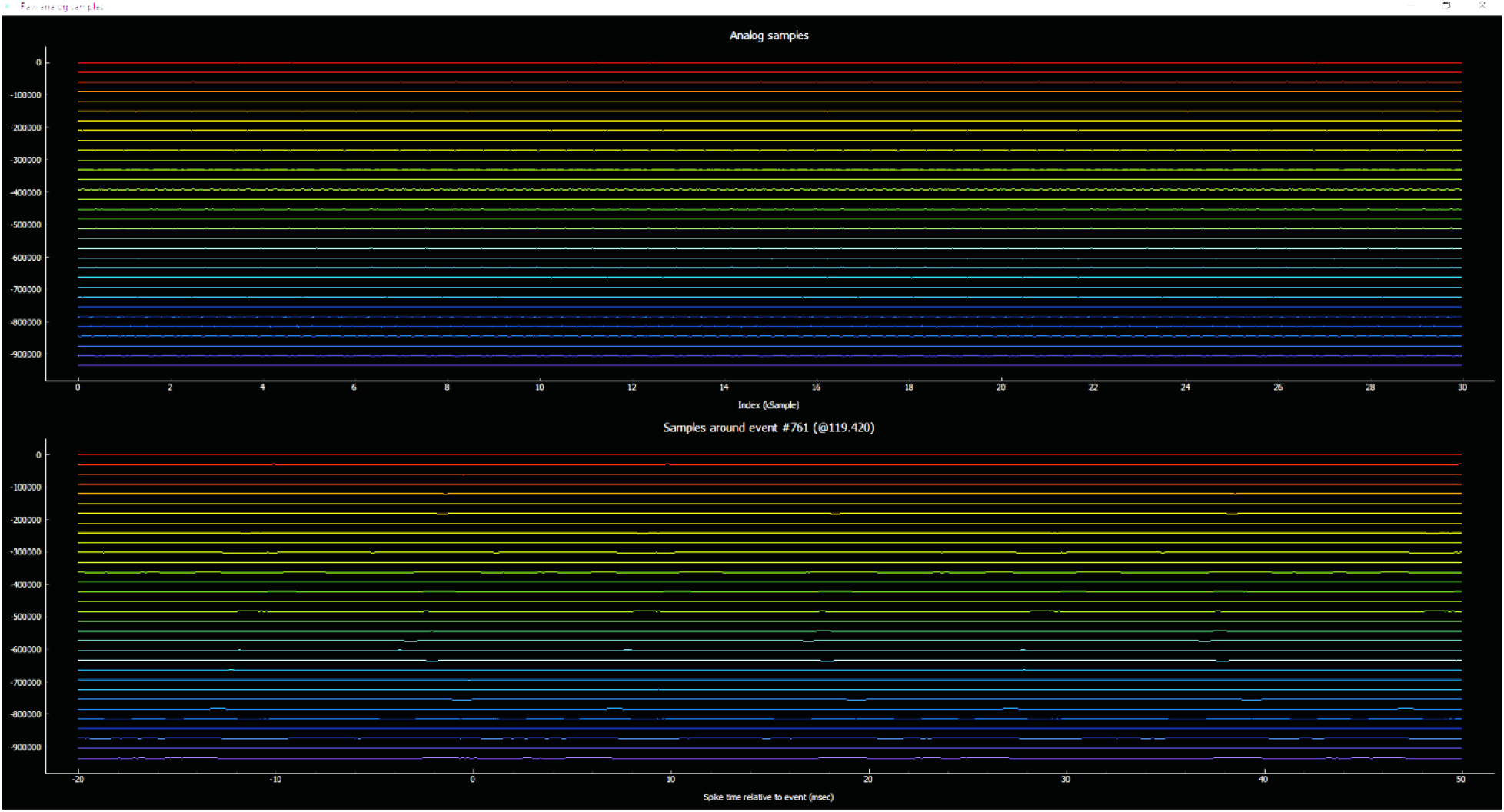
Raw analog data window

